# Disentangling the origins of confidence in speeded perceptual judgments through multimodal imaging

**DOI:** 10.1101/496877

**Authors:** Michael Pereira, Nathan Faivre, Iñaki Iturrate, Marco Wirthlin, Luana Serafini, Stéphanie Martin, Arnaud Desvachez, Olaf Blanke, Dimitri Van De Ville, José del R. Millán

**Affiliations:** Defitech Foundation Chair in Brain-Machine Interface, École Polytechnique Fédérale de Lausanne, Geneva, Switzerland; Center for Neuroprosthetics, École Polytechnique Fédérale de Lausanne, Geneva, Switzerland; Laboratory of Cognitive Neuroscience, Brain Mind Institute, Faculty of Life Sciences, École Polytechnique Fédérale de Lausanne, Geneva, Switzerland; Centre d’Economie de la Sorbonne, CNRS UMR 8174, Paris, France; Department of Biomedical, Metabolic and Neural Sciences, University of Modena and Reggio Emilia, Modena, Italy; Department of Neurology, University Hospital Geneva, Geneva, Switzerland; Medical Image Processing Lab, Institute of Bioengineering, École Polytechnique Fédérale de Lausanne, Geneva, Switzerland; Department of Radiology and Medical Informatics, University of Geneva, Geneva, Switzerland

**Keywords:** metacognition, error monitoring, confidence, EEG, fMRI, race modeling, inferior frontal gyrus, insula, anterior prefrontal cortex

## Abstract

The human capacity to compute the likelihood that a decision is correct - known as metacognition - has proven difficult to study in isolation as it usually co-occurs with decision-making. Here, we isolated post-decisional from decisional contributions to metacognition by combining a novel paradigm with multimodal imaging. Healthy volunteers reported their confidence in the accuracy of decisions they made or decisions they observed. We found better metacognitive performance for committed vs. observed decisions, indicating that committing to a decision informs confidence. Relying on concurrent electroencephalography and hemodynamic recordings, we found a common correlate of confidence following committed and observed decisions in the inferior frontal gyrus, and a dissociation in the anterior prefrontal cortex and anterior insula. We discuss these results in light of decisional and post-decisional accounts of confidence, and propose a generative model of confidence in which metacognitive performance naturally improves when evidence accumulation is constrained upon committing a decision.

## Introduction

Upon making decisions, one usually “feels” that a given choice was correct or not, which allows deciding whether to commit to the choice, to seek more evidence under uncertainty, or to change one’s mind and go for another option. This crucial aspect of decision making relies on the capacity to monitor and report one’s own mental states, which is commonly referred to as metacognitive monitoring (Fleming et al., 2012; Koriat, 2006). One promising venue to unravel the neural and cognitive mechanisms of metacognitive monitoring involves investigating how, and to what extent, humans become aware of their own errors (Yeung & Summerfield, 2012). Typically, volunteers are asked to execute a first-order task under time pressure (e.g., numerosity: which of two visual arrays contains more dots) and afterward perform a second-order task by providing an estimate of confidence in their response (“how sure were you that your response was correct?”). Confidence is formally defined as the probability that a first-order response was correct given the available evidence (Pouget et al., 2016). Distinct models have been proposed to explain how confidence is computed: some models consider confidence as a fine-grained description of the same perceptual evidence leading to the first-order decision (Kiani & Shadlen, 2009), sometimes enriched with post-decisional processes (Pleskac et al., 2010, Van Den Berg et al., 2016; Fleming et al., 2017). Other models posit that confidence stems from mechanisms different from those responsible for making that decision (for review, see Grimaldi et al., 2015). However, as of today, the contribution of (post-)decisional signals on confidence remains unclear, principally due to the difficulty of dissociating confidence from first-order decision-making.

Here we combined a novel paradigm with multimodal neuroimaging to dissociate confidence from decision-making. Our paradigm allowed a controlled comparison of confidence ratings for decisions that were *committed* (i.e., taken and reported by participants), and decisions that were merely *observed* (i.e., taken by a computer). Hereby, we could isolate the contribution of decisional signals to confidence (Figure 1A). In the *active* condition, 20 participants were presented with two arrays of dots for 60 ms and were asked to indicate which of the two arrays contained more dots by pressing a button with the left or right hand (numerosity first-order task). At the end of each trial, participants had to report their confidence in their response being correct or incorrect using their left hand (second-order task). The *observation* condition followed the exact same procedure, except that the first-order task was performed automatically: participants saw the image of a hand over the right or left array of dots with identical yet shuffled timings and choice accuracy (i.e., observation trials were a shuffled replay of active trials, see methods). They were then asked to report their confidence in the observed decision. This allowed us to quantify metacognition for committed (active condition) compared to observed (observation condition) decisions while keeping perceptual evidence, first-order performance, and timing constant across conditions. Both conditions were performed while recording simultaneous electroencephalography (EEG) and functional magnetic resonance imaging (fMRI), to constrain blood-oxygenation level dependent (BOLD) correlates of confidence to electrophysiological processes occurring immediately after the committed or observed decision.

**Figure 1.**
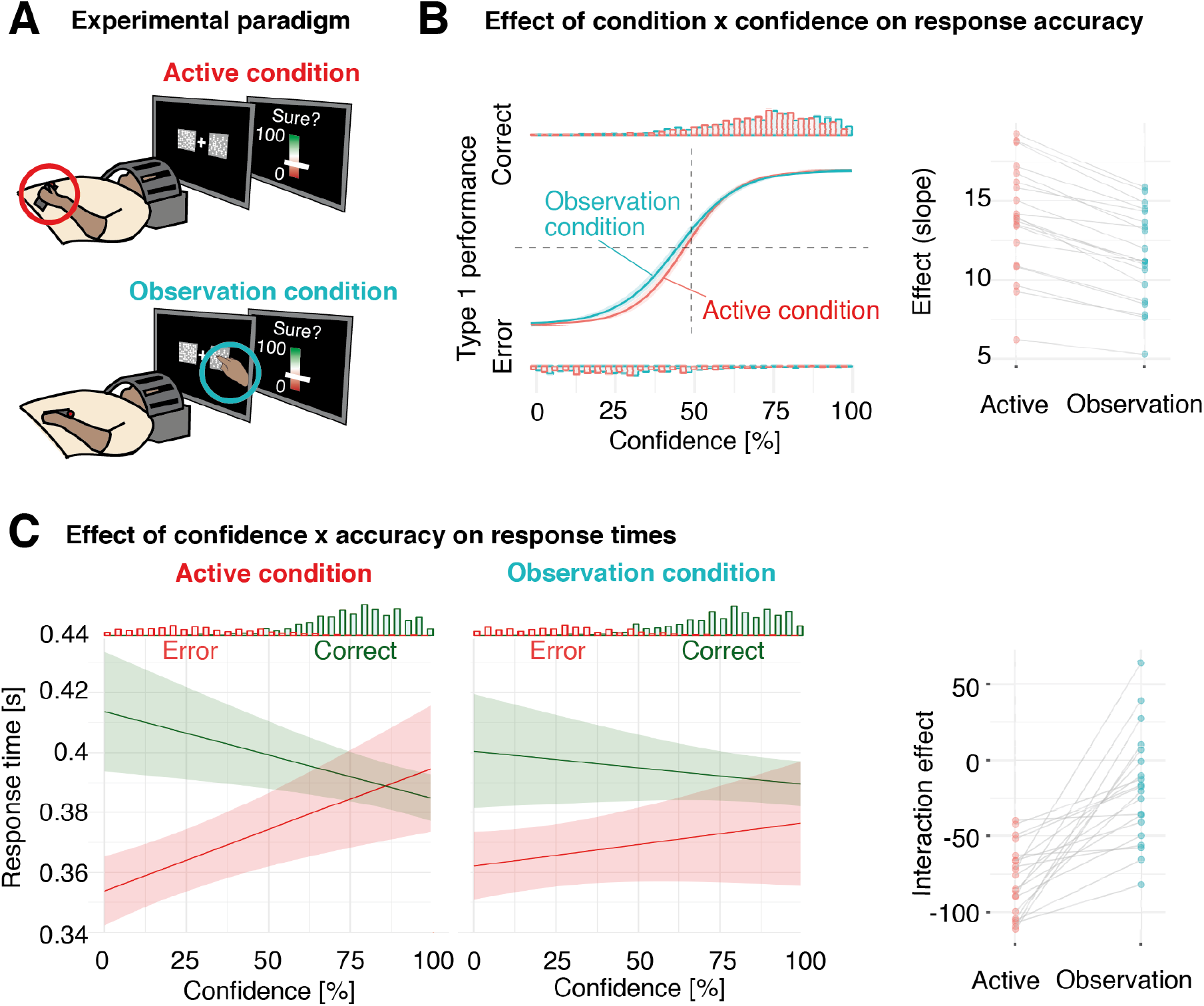
Experimental paradigm and behavioral results. (A) Experimental paradigm: a participant lying in the fMRI bore equipped with an EEG cap performs (active condition in red) or observes (observation condition in blue) the first-order task, and subsequently reports confidence in the committed or observed decision using a visual analog scale. (B) Mixed effects logistic regression between first-order accuracy and confidence in the active (red) and observation condition (blue). The histograms represent the distributions of confidence for correct (top) and incorrect (bottom) first-order responses. Right panel: Individual slopes of the mixed effects logistic regression indicating metacognitive performance. (C) Mixed effects linear regression between first-order response times and confidence for correct (in green) and incorrect trials (in red) in the active (left panel) and observation condition (right panel). The histograms represent the distributions of response times and confidence for correct and incorrect first-order responses. Rightmost panel: interaction term between first-order accuracy and confidence for response times in the active compared to observation condition. Shaded areas represent 95% confidence intervals.

Data collection was conducted in view of testing three pre-registered hypotheses (https://osf.io/a5qmv). At the behavioral level, assuming that signals associated with overt decisions inform confidence judgments, we expected confidence ratings to better track first-order performance for committed compared to observed decisions. Based on several findings showing a role of action monitoring for confidence (e.g., Fleming & Daw, 2017; Fleming et al., 2015; Faivre et al., 2018), we expected brain regions encoding confidence specifically for committed decisions to be related to the cortical network involved in action monitoring, and brain regions conjunctively activated in both conditions to reflect a shared mechanism independent from decision commitment. Finally, we expected to find earlier correlates of confidence following committed compared to observed responses, as efferent information is available before visual information (Holroyd and Coles, 2002).

## Results

### Better metacognitive performance for committed compared to observed decisions

The influence of decision commitment on second-order judgments was assessed by comparing metacognitive performance for committed compared to observed decisions. The first-order task consisted of indicating which of two arrays contained more dots (active condition), or observing a hand making that decision (observation condition) (Figure 1A). By design, first-order performance was identical in the two conditions (see Methods), with an average first-order accuracy of 71.2 % (± 1.0 %, 95 % CI, according to a 1up/2down adaptive procedure), first-order response time of 385 ms ± 8 ms, and difference of 13.1 ± 1.7 dots between the two arrays.

We then turned to second-order performance, quantifying metacognitive performance as the capacity to adapt confidence to first-order accuracy. Confidence was measured on a continuous scale quantifying the probability of being correct or incorrect (ranging from 0: “sure error” to 1: “sure correct”). A mixed effects logistic regression on first-order accuracy as a function of confidence and condition revealed an interaction between confidence and condition (model slope: odds ratios z = 2.90, p = 0.004; marginal R^2^ = 0.69), indicating that the slope between confidence and first-order accuracy was steeper in the active compared to observation condition (Figure 1B). This difference in metacognitive performance was present in all participants we tested, and also found when analyzing the data with tools derived from second-order signal detection theory (area under the type II receiver operating curve (AROC): active condition = 0.92 ± 0.02; observation condition = 0.90 ± 0.03; Wilcoxon sign rank test: V = 163, p = 0.03, see SI). In addition, metacognitive performance was correlated between conditions (R2 = 0.93, p < 0.001), suggesting partially overlapping mechanisms for monitoring committed and observed decisions. Of note, confidence per se did not differ across conditions (F(1,4772) = 0.01, p = 0.98).

To assess the contribution of decisional signals to metacognitive monitoring, we ran a linear mixed effects model on first-order response times as a function of confidence, accuracy, and condition. This model revealed a triple interaction (F(1,4742) = 6.05, p = 0.014), underscoring that in the active condition, response times for correct responses correlated negatively with confidence, and response times for errors correlated positively with confidence (F(1,26) = 23.70, p < 0.001, Figure 1C). No main effect of confidence (F(1,29) = 0.02, p = 0.89) nor interaction between confidence and accuracy (F(1,19) = 1.34, p = 0.26) was observed in the observation condition (Figure 1C). Together, these results indicate that confidence was modulated by committed but not observed response times, and thus suggest the importance of decisional signals and potentially motor actions to build accurate confidence estimates.

To further elucidate the contribution of response times to confidence, we ran follow-up experiments including a third condition in which the first and second-order responses were reported simultaneously on a unique scale. We were able to replicate our finding of higher metacognitive performance between the active and observation condition, and found that metacognitive performance in the active condition was better than when first and second-order responses were provided simultaneously. This confirms that the readout of speeded motor actions associated with decision commitment serves subsequently as input to compute confidence. Lastly, to rule out the possibility that increases in metacognitive performance were due to confounding factors between the active and observed conditions (e.g., demand characteristics, visual saliency), we performed the same experiment under no-time pressure, and found no difference in metacognitive performance between committed and observed decisions (see SI). Altogether, these results validate our first pre-registered hypothesis that metacognitive performance is better for committed compared to observed speeded decisions, and suggest that action monitoring might play a role in this process.

### BOLD correlates of confidence

We sought to find the brain regions co-activating with confidence by parametrically modulating a general linear model (GLM) with participants’ confidence ratings, as well as response times and perceptual evidence (i.e., the difference in number of dots between the right and left side of the screen) as regressors of no interest (see methods). Because error monitoring and confidence are tightly related (Yeung & Summerfield, 2012), we deliberately analyzed the neural correlates of confidence without modeling first-order accuracy. Of note, the visual scale we used allowed participants to report their confidence estimate with a single and identical motor action with the left hand across conditions and trials, ruling out motor confounds when analyzing data (see methods). Widespread activity correlating both positively and negatively with confidence was found in the active and observation condition, in line with several other studies (Fleck et al., 2005; Fleming et al., 2012b; Baird et al., 2013; Heereman 2015; Hebart et al., 2016; Morales et al., 2018; Vaccaro & Fleming, 2018). A complete list of activations can be found in Supplementary Table 1. In addition, we found that the right precentral gyrus (contralateral to the hand reporting confidence), left insula, and bilateral pMFC were significantly more predictive of confidence in the active than in the observation condition (Supplementary table 2). We then defined the regions commonly activated by confidence in both conditions. A conjunction analysis revealed that the bilateral pMFC, left IPL, precentral gyrus, AI and IFG were negatively correlated with confidence (Figure 2B; Supplementary Table 3).

**Figure 2.**
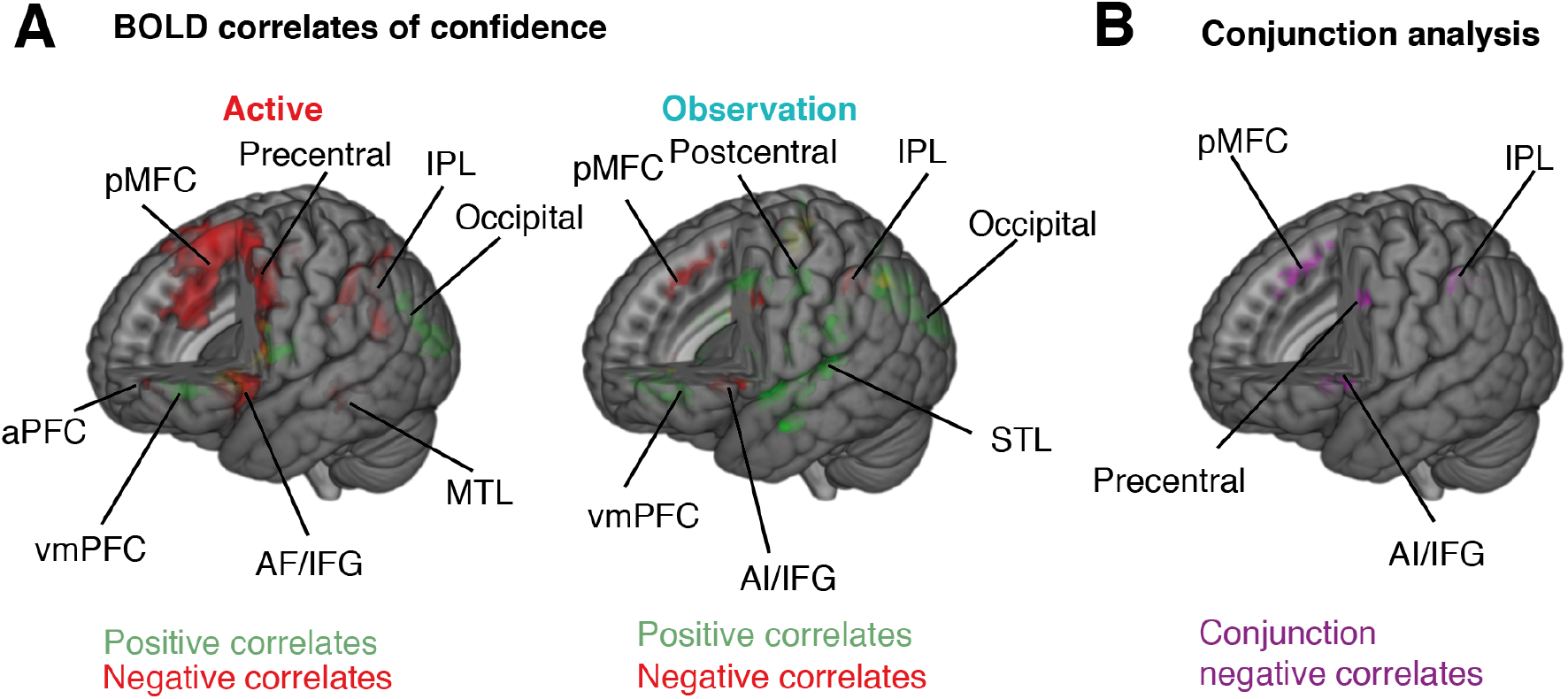
BOLD correlates of confidence. (A) Brain areas co-activated with positive (green) and negative (red) confidence values for the active (left) and observation (right) conditions. (B) Brain areas co-activated with negative confidence values in both conditions (conjunction analysis). All displayed BOLD activations are FWE-corrected (p<0.05) at the cluster-level with a threshold at p<0.001. Labels: anterior insula (AI), anterior prefrontal cortex (aPFC), Posterior medial frontal cortex (pMFC), inferior frontal gyrus (IFG), inferior parietal lobule (IPL), medial temporal lobe (MTL), superior temporal lobe (STL), ventromedial prefrontal cortex (vmPFC). Not all brain regions are labeled (see Supplementary Tables 1-3).

#### ERP correlates of confidence

To further isolate the neural correlates of confidence for committed and observed decisions, we identified which regions co-activated with EEG correlates of confidence occurring exclusively within five hundred milliseconds after the first-order response (i.e., post-decisional processes). We first modeled the EEG amplitude time-locked to the first-order response as a function of confidence using mixed effects linear regression, with first-order response times and perceptual evidence as covariates of no interest (see methods). In the active condition, we found that EEG amplitude correlated with confidence starting 68 ms following the first-order response over centro-parietal electrodes, resembling a centro-parietal positivity (CPP; Figure 3A, top left; O’Connell et al., 2012). Another correlate of confidence was found 88 ms post-response over frontoparietal electrodes, akin to an error-related negativity (ERN; Figure 3A, bottom left; Falkenstein et al., 1991, Gehring et al., 1993). In the observation condition, correlates of confidence were found on the same two electrodes with similar topography (correlation between frontocentral cluster in the active and observation conditions: rho = 0.88) but not before 200 ms post-response (Figure 3A, right).

**Figure 3.**
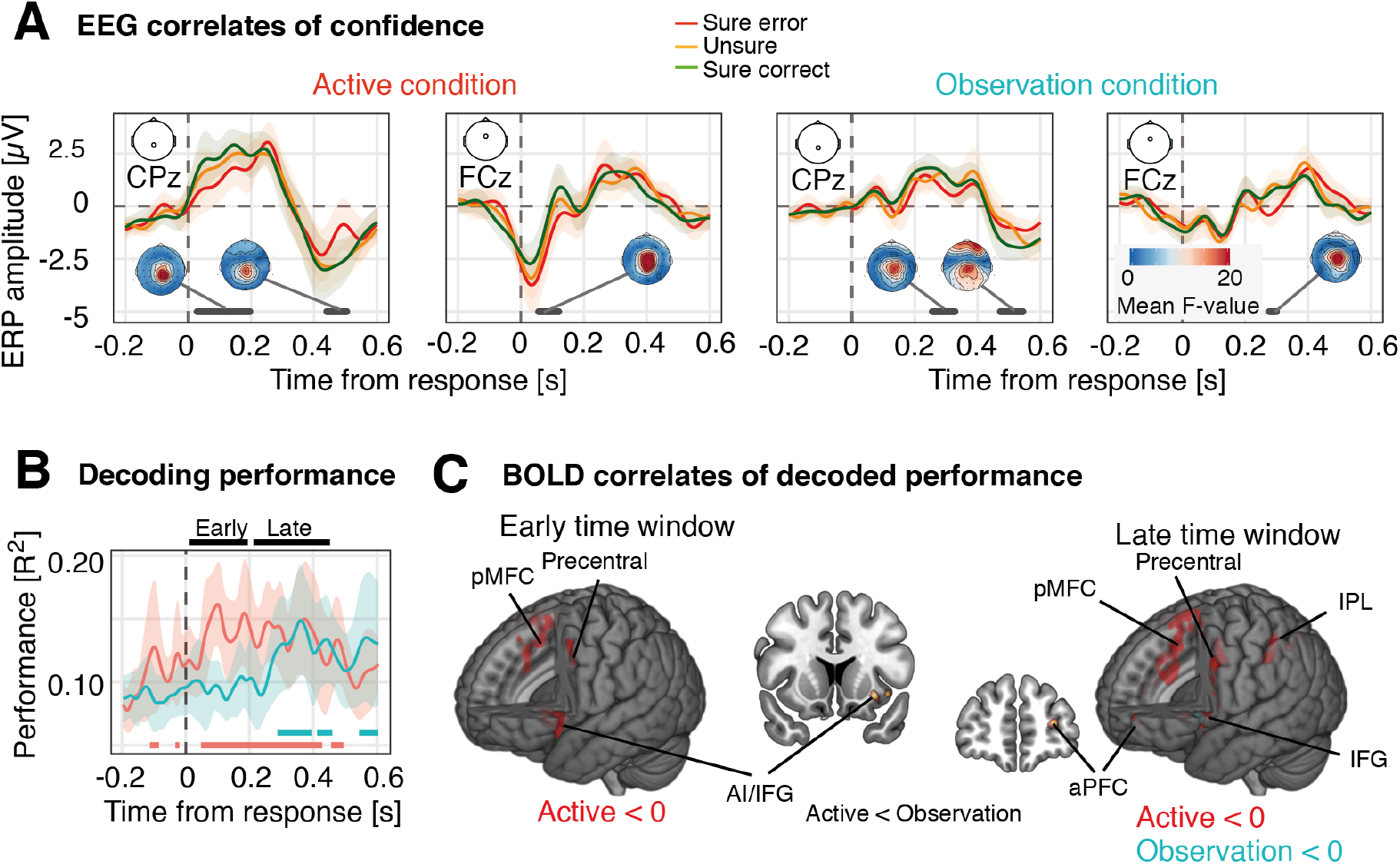
EEG-informed correlates of confidence. (A) ERPs time-locked to the first-order response are shown for the active condition (left panels) and observation condition (right panels) for the CPz and FCz sensors. For illustrative purposes, epochs were binned according to three levels of reported confidence: sure error (0 - 33% confidence), unsure (34 - 66% confidence) and sure correct (67 - 100% confidence), although statistics were computed with raw confidence values using mixed effects linear regression. The shaded areas represent 95%-CI. Regions of significance (p<0.05, FWE corrected) are depicted with a gray line, along with topographic maps of the corresponding F values. (B) Leave-one-out decoding performance over time. The plot shows the amount of variance of the reported confidence explained by the decoder (R^2^) over time in the active (red trace) and the observation condition (blue trace). The shaded areas represent 95%-CI, and the horizontal dashed lines the chance level (p<0.05, computed via non-parametric permutation tests corrected for multiple comparisons). For each participant and condition, the output of the best decoder within an early and late time window was retrained on the whole dataset and used as a parametric regressor to model the BOLD signal. (C) Brain areas co-activated with low decoded-confidence values in the early (left) and late time window (right). All displayed BOLD activations are FWE-corrected (p<0.05) at the cluster-level with a threshold at p<0.001. Labels: Posterior medial frontal cortex (pMFC), inferior parietal lobule (IPL), anterior insula (AI), inferior frontal gyrus (IFG) and anterior prefrontal cortex (aPFC). Not all brain regions are labeled (see Supplementary Table 4). The coronal view shows significant differences between the active and the observation condition for the labelled region (AI for the early time window and aPFC for the late time window).

### Common and distinct BOLD correlates of EEG decoded confidence

The brain regions corresponding to the ERP correlates of confidence were identified by modeling the BOLD signal with EEG-based single-trial predictions of confidence. Confidence predictions at each time point were derived from a linear regressor taking the EEG independent components activation profiles as low-dimensional variables (N=8 ± 3 for each participant, see methods). Leave-one-out performance was significant at the group level (non-parametric permutation test, corrected p < 0.05) with a peak decoding performance achieved 96 ms and 356 ms following committed and observed responses (Figure 3B).

To dissociate early correlates of potentially “all-or-none”, binary error detection from fine-grained second-order confidence estimates described as occurring 200 ms after response (Boldt and Yeung, 2015), we selected two time points corresponding to local peaks in the cross-validated decoding performance within an early (50 - 200 ms post response) and late (200 - 450 ms) temporal windows (see Methods). The latency of the early peaks was 108 ± 22ms in the active condition. There was no significant decoding in the early time window in the observation condition. Late peak latencies were 321 ± 31 ms in the active and 353 ± 27 ms in the observation condition, with no significant difference between condition (t(19) = - 1.49, p = 0.15). Based on these two time-points, we re-trained one regressor per condition and peak on all available epochs and used the resulting single-trial predictions as a parametric regressor to model the BOLD signal, along with first-order response times and perceptual evidence as covariates of no interest. By using EEG as a time-resolving proxy to BOLD signal (Britz et al., 2010), we sought to investigate the anatomical correlates of confidence at specific timings, with the aim of disentangling BOLD signal associated with pre and post-decisional processes (Gherman & Philiastides 2018).

The regions co-activating with *decoded* confidence in the early time window included the bilateral pMFC, the left IFG, AI and MFG (Figure 3C, left). For the late time window (Figure 3C, right), coactivations with low decoded-confidence were found in the bilateral pMFC and IFG, the left precentral gyrus, IPL, AI, MFG and aPFC for the active condition, and in the left IFG for the observation conditions (Supplementary Table 4). The left IFG was thus commonly activated by low decoded-confidence in both conditions. Differences between co activations in the active and observation condition were found in the anterior insula (AI) in the early time window and in the aPFC in the late time window (Figure 3C; Supplementary Table 4).

### Behavioral modeling

In view of obtaining a mechanistic understanding of the way decisional and post-decisional evidence contribute to confidence, we derived confidence in committed and observed decisions using a race accumulator model, considered to be biologically plausible representations of evidence accumulation in the brain (Bogacz et al., 2006; Gold and Shadlen, 2007). Such models assume that ideal observers commit to a first-order decision (D; Figure 4A) once one of two competing evidence accumulation processes (here, corresponding to evidence for the left or right choice) reaches a decision.

**Figure 4.**
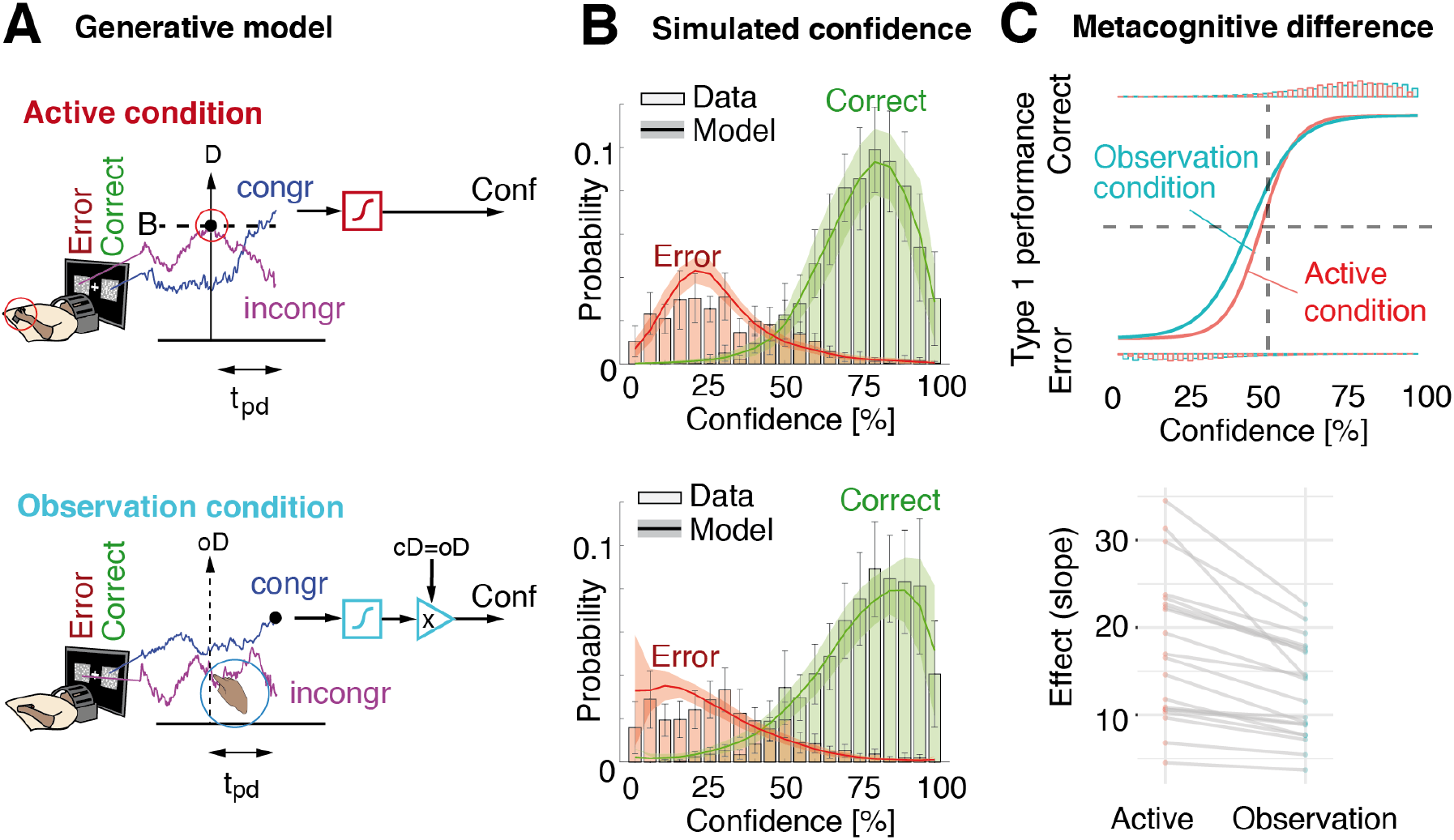
Race accumulator model for confidence. (A) Upper plot: an example trial for which the participant made a first-order error. The violet and blue traces represent accumulators that are incongruent and congruent with a correct response, respectively. A committed first-order decision (D) is taken when the winning accumulator hits the decision bound (dashed horizontal line). Here, the violet trace wins, producing a first-order error. Confidence is assumed to be based on the difference between both accumulators at the end of the post-decisional period. Similarly, confidence in the observed response is read-out from the difference between both accumulators at the end of the post-decisional period. In both plots, the sigmoid (square box) constrains the result to the [0,100] % interval. T_nd is the non-decisional time, t_d the time taken for the winning accumulator to reach the decision bound B and t_pd the post-decisional time. (B) Histogram of the confidence ratings obtained during the experiments, compared to the model simulations (thick line) for error (red) and correct (green). Upper plot for the active condition (second-order model), lower plot for the observation condition (non-decisional model). Error bars and shaded area represent 95% confidence intervals across subjects. (C) Top panel: Mixed logistic regression between simulated first-order accuracy and simulated confidence, in the active (red) and observation condition (blue). Bottom panel: Individual slopes of the mixed regression model indicating metacognitive performance, see Figure 1B for the actual behavioral results.

We first fitted five parameters (i.e., drift, bound, non-decision time, non-decision time variability and starting point variability, see methods) to first-order choice accuracy and response times recorded for each participant during the active condition. With these parameters, we simulated pairs of competing evidence accumulation trajectories leading to first-order choices and response times. We then derived confidence based on a mapping of the state of evidence of the winning accumulator, following recent findings that confidence is based solely on evidence supporting the decision (Peters et al. 2018; Zylberberg et al., 2012). To account for changes-of-mind, we sampled accumulated evidence after a post-decisional period (tpd in Figure 4A; Peskac and Busemeyer, 2010; Van Den Berg et al., 2016) corresponding to the average peak decoding accuracy found with EEG (see previous section). The sampled evidence was mapped to the range of confidence ratings using a sigmoidal transformation with two additional free parameters controlling for bias and sensitivity (see methods).

For the observation condition, we assumed a similar evidence accumulation process, except that choice and response times were independent from the evidence accumulation process, as in our paradigm. Since first-order behavior in the observation condition remained latent by design, we used the parameters fitted for the active condition to simulate a second dataset of pairs of competing evidence accumulation trajectories. We then mapped confidence from a readout of the accumulator with highest evidence after the post-decisional period, but time-locked to shuffled observed decisions (oD in Figure 4A) and response times, as in our paradigm. When observed decisions were incongruent with covert decisions, we inverted the simulated confidence ratings. This model fitted confidence data better than an alternative model for which participants did not make covert decisions and simply readout confidence from the state of evidence of the accumulator corresponding to the computer’s choice (log-likelihood: −2.13 ± 6.32 versus −2.91 ± 6.65, Wilcoxon sign rank test, p = 0.019).

Across participants, our model fitted confidence ratings well (active condition: R2 = 0.71 ± 0.30; observation: R2 = 0.65 ± 0.40; Figure 4B), suggesting that it represents a plausible mechanism of confidence build-up for speeded decisions. Most importantly, the confidence model for the active condition predicted better metacognitive accuracy than the observation model, consistent with our experimental data (Figure 4C). As in the behavioral analysis, we ran a mixed effects logistic regression on first-order accuracy as a function of confidence and condition, which revealed an interaction between confidence and condition (odds ratios z = −4.58, p < 0.001), indicating that the slope between confidence and first-order accuracy was steeper in the active compared to observation condition. Area under the type II receiver operating curve (AROC) was also higher for the active condition (0.95 ± 0.02 vs. 0.93 ± 0.03, Wilcoxon sign rank test, V = 197, p < 0.001). Of note, these differences were not explained by differences in the goodness-of-fit across subjects (R=0.13; p=0.59). We could thus reproduce the lower metacognitive performance found in the observation condition only by detaching the decision process from the evidence accumulation process leading to confidence.

## Discussion

The present study evaluated the contribution of decisional signals to metacognition by comparing and modeling confidence judgments for committed and observed decisions, and identifying the neural correlates of confidence with high spatiotemporal resolution. A group of 20 healthy volunteers was asked to perform or observe a perceptual task, and then indicate their confidence regarding the accuracy of the committed or observed decisions.

### Better metacognitive performance for committed decisions

Participants were able to adjust confidence to the accuracy of their own perceptual decisions, and to the accuracy of decisions they observed. Yet, consistent with our pre-registered predictions, committed decisions were associated with a slight but consistent increase in metacognitive performance compared to observed decisions, which supports decision commitment as an additional input for confidence. Of note, this effect could not be explained by differences in terms of perceptual evidence or first-order performance across conditions, which were identical by design (see Methods). A follow-up experiment revealed equivalent metacognitive performance for committed and observed decisions when participants were given more time to perform the first-order task. This indicates that the metacognitive advantage we describe occurred in speeded tasks in which errors are immediately recognized as such (Charles et al., 2013). By showing the specificity of metacognitive improvement for committed decisions under speeded conditions, this follow-up experiment also undermines the possibility that our effect stems from experimental confounds between the active and observation conditions (e.g., demand characteristics, visual saliency), as such confound would likely pertain both to speeded and non-speeded conditions. Last, we found that metacognitive performance in the active condition was better than another condition involving simultaneous first and second-order responses, in which by definition confidence could not be informed by a previous committed decision. This brings another line of evidence that action monitoring plays a role for confidence.

We then turned to computational modeling to shed light on the role of decisional signals for decision monitoring (Kepecs et al., 2008, Kiani et al., 2009, Pleskac et al., 2010, Maniscalco & Lau, 2016). One biologically plausible (computational account of decision making, called race accumulator model (Bogacz et al., 2006; Kiani et al., 2014), assumes that ideal observers commit to a first-order decision (here, the right or left side of the screen containing more dots) once one of two competing evidence accumulation processes (for one or the other choice) reaches a decision boundary. We extended these models, assuming a continuation of evidence accumulation after the first-order decision (Van Den Berg et al., 2016). Through this procedure, we found that the path of second-order evidence accumulation in the active condition was constrained by the first-order decision boundary, which translated into confidence estimates with lower variance compared to observed responses which impose no constraint on evidence accumulation (7.24 ± 0.11 vs 9.04 ± 0.16, Wilcoxon signed rank test, V = 8, p < 0.001). This prediction was verified a posteriori in our behavioral data, as we found higher variance for confidence ratings in the observation vs. active condition (6.71 % ± 0.92 vs. 7.33 ± 1.15, Wilcoxon signed rank test, V = 45, p = 0.024).

The notion that committing to (but not observing) first-order decisions sharpens confidence estimates is corroborated by studies showing that metacognitive performance increases when response times are taken into account to compute confidence (Siedlecka, Paulewicz, & Wierzchoń, 2016), and decreases in case motor actions are irrelevant to the task at play (Kvam et al., 2015), or when the task-relevant motor action is disrupted by transcranial magnetic stimulation over premotor cortex (Fleming et al., 2015). The role of motor signals for metacognition is also supported by recent results indicating that confidence increases in presence of sub-threshold motor activity prior to first-order responses (Gajdos et al., 2018); and that alpha desynchronization over the sensorimotor cortex controlling the hand performing that action correlate with confidence (Faivre et al., 2018). Together, these empirical results suggest that confidence is not solely derived from the quality of perceptual evidence, but involves the perception-action cycle. By comparing committed and observed decisions in a controlled way, we could test a direct prediction derived from these studies, and document its neural and computational mechanisms.

### Neural correlates of confidence in committed and observed decisions

After assessing the contribution of decision commitment to confidence at the behavioral level, we identified the brain regions at play for monitoring committed and observed decisions by parametrically modulating the BOLD signal by confidence estimates. Besides brain regions activated independently across conditions (Supplementary table 1), we found that the right precentral gyrus (contralateral to the hand reporting confidence), left anterior insula and bilateral pMFC were significantly more predictive of confidence in the active than in the observation condition (Supplementary table 2). The involvement of such motor and error detection regions (Carter et al., 1998; Bonini et al., 2014; Bastin et al., 2017), together with our behavioral and modeling results support the notion that action monitoring serves as input for confidence. This is corroborated by behavioral results from a follow-up experiment, showing that metacognitive performance was better in the active condition compared to a condition in which the first and second-order responses were reported simultaneously on a unique scale.

In search for hemodynamic correlates of confidence independent from action commitment, we identified the brain regions conjunctively related to confidence in the active and observation conditions as the pMFC, insula, IFG, IPL and precentral gyrus (See Supplementary Table 3). This is corroborated by previous results by Heereman and colleagues (2015), who found the pMFC, insula and IFG to be negatively correlated with confidence during motion and color discrimination tasks, as well as Morales and colleagues (2018), who found the pMFC to be negatively correlated for confidence in perceptual and memory tasks. In addition, IPL activations (Hayes et al., 2011; Kim & Cabeza, 2007, 2009; Moritz et al., 2006) and gray matter thickness (Filevich et al., 2018) were shown to correlate negatively with confidence. These regions could represent a substrate for the computation of confidence, stripped from decisional and error correction processes.

### Timing of confidence-related brain activations

Due to the low temporal resolution of the BOLD signal, it is worth considering that the above-mentioned regions may be contaminated by prerequisites of confidence computation (e.g., quality of numerosity representation, alertness), as well as its by-products (e.g., the act of reporting confidence on the scale). To further isolate the neural correlates at play when computing confidence for committed and observed decisions and pruning out some of the prerequisites and by-products of confidence, we constrained our search to neural events occurring in the vicinity of the committed/observed first-order response by fusing EEG and fMRI data (Debener et al., 2005; Gherman & Philiastides, 2018).

In line with our pre-registered hypothesis, we found early correlates of confidence for committed but not for observed decisions in fronto-central EEG activity resembling the error-related negativity (ERN) involved in error detection (Boldt & Yeung, 2015) and in fronto-parietal activity resembling the centro-parietal positivity (CPP) involved in evidence accumulation (O’Connell et al., 2012). To address the possibility that early correlates of confidence in observed decisions do not appear in event-related potentials but involve multivariate electrophysiological patterns, we built a decoder of confidence based on whole-scalp EEG. Coherently with the univariate results described above, our decoder could explain confidence better than chance level in the time vicinity of committed decisions (108 ms post-response), while significant decoding performance was only attained 353 ms after observed decisions. The absence of early correlates of confidence in the observation condition was expected as participants could not possibly assess first-order accuracy before perceiving the observed decision (Holroyd & Coles 2002, Van Schie et al., 2004; Iturrate et al., 2015). Of note, decoding performance in the active condition plateaued after the first peak and dropped after around 400 ms, indicating that ongoing processes leading to confidence may be sustained in time. Thus, the computation of confidence may unfold in two waves, an early one specific to the the monitoring of committed decisions, and a later one for computing confidence *per se*. One possibility is that the early correlate for committed decisions relates to an “all-or-none” automatic error detection system (Charles et al., 2013, although see Vocat et al., 2011, Pereira et al., 2017), while the late correlate underlies a fine-grained estimation of second-order signals (Boldt & Yeung, 2015).

We finally examined the properties of early and late correlates of confidence by assessing their BOLD covariates. For that, we parametrically modulated the BOLD signal using the output of a decoding model of confidence based on whole-scalp EEG, hereby obtaining a time-resolved description of fMRI data (Gherman & Philiastides, 2018). In the active condition, we found that the pMFC, IFG, MFG and insula were co-activated both during the early and late decoding window. These regions are likely to relate to early error processing based on the monitoring of errors/conflicts surrounding the first-order response (Dehaene et al., 1994, Carter et al., 1998, Bonini et al., 2014, Bastin et al., 2017, Ullsperger et al., 2014 for a review). Furthermore, Murphy and colleagues showed that similar error-related feedback signals from the pMFC inform metacognitive judgments through the modulation of parietal activity involved in evidence accumulation (Murphy et al., 2015). Other regions including the IPL, precentral cortex and aPFC were found specifically in the late decoding window, which hints to their involvement in late processes at play for the computation of graded confidence estimates. In the observation condition, the only region coactivated with late electrophysiological correlates of confidence was the left IFG, adjacent to the cluster we found in the active condition. This suggests the role of left IFG operating similarly around 300 ms whether a decision is committed or observed. Of note, the quest for domain-general mechanisms of confidence (Faivre et al., 2018, Rouault et al., 2018) is hindered by the fact that our paradigm alternated short blocks of active and observation conditions, which could potentially inflate correlations in confidence due to confidence leaks across trials (Rahnev et al., 2015).

By contrast to decision-independent activations in the IFG, the aPFC – commonly referred to as a key region for confidence (Fleming et al., 2010, 2012, Morales et al., 2018, for review see Grimaldi et al., 2015)– was involved in monitoring committed decisions only. The fact that activity in the insula and aPFC were not related to confidence in observed decisions reveals that these regions may underlie a putative role in linking first-order decisional signals allowing early error detection to inform fine-graded confidence estimates derived from the quality of perceptual evidence (Fleming et al., 2018). Beyond error detection, the aPFC could operate by linking other sources of information to inform confidence, including the history of confidence estimates over past trials (Shekhar et al. 2018). Although this claim deserves further investigations, it extends a recent proposal by Bang & Fleming (2018) arguing that aPFC is involved in reporting rather than computing confidence estimates perse.

## Conclusion

We combined psychophysics, multimodal brain imaging, and computational modeling to unravel the mechanisms at play when monitoring the quality of decisions we make, in comparison to equivalent decisions we observe. Our behavioral and modeling results indicate that committing to a decision leads to increases in metacognitive performance, presumably due to the constraint of evidence accumulation by first-order decisions. By focusing the analysis of neural signals on processes independent from decision-making, we isolated the IFG as a key region contributing to confidence in both committed and observed decisions. We further specified the functional role of the IFG, distinct from a set of regions involved in error processing, and from the insula and aPFC which could potentially inform confidence estimates with the output of such error processing.

## Supporting information

## Methods

Software and algorithms

| Reagent or resource | Source | Identifier |
| --- | --- | --- |
| MATLAB 2017a | Mathworks<br>http://www.mathworks.com/products/matlab/ | **RRID:SCR_001622:** |
| SPM12 | https://www.fil.ion.ucl.ac.uk/spm/software/spm12/ | **RRID:SCR_007037** |
| EEGLAB | http://sccn.ucsd.edu/eeglab/index.html | **RRID:SCR_007292** |
| Analyzer | BrainVision | **RRID:SCR_002356** |
| R | http://www.r-project.org/ | **RRID:SCR_001905** |
| ggplot2 | http://ggplot2.org/ | **RRID:SCR_014601** |
| lme4 | https://cran.r-project.org/web/packages/lme4/index.html | **RRID:SCR_015654** |

## CODE AVAILABILITY

Matlab and R code for reproducing all analyses can be found on GitHub (https://gitlab.com/nfaivre/analysis_public).

## DATA AVAILABILITY

All data, analysis and modeling software scripts from this study will be made freely available upon publication. Anonymized data will be stored on openneuro.org. Unthresholded statistical maps can be found on NeuroVault (https://neurovault.org/collections/4676/)

## EXPERIMENTAL MODEL AND SUBJECT DETAILS

The experimental paradigm, sample size, and analysis plan detailed below were registered prior to data collection using the open science framework (https://osf.io/a5qmv).

Twenty-five healthy volunteers (12 females, mean age = 24.6 ± 1.43) from the student population at the Swiss Federal Institute of Technology took part in this study in exchange for monetary compensation (20 CHF per hour). All participants were right-handed, had normal hearing and normal or corrected-to-normal vision, and no psychiatric or neurological history. They were naive to the purpose of the study and gave informed consent. The study was approved by the ethical committee of the canton of Geneva, Switzerland (Commission Cantonale d’Ethique de la Recherche (CCER); study number 2017-00014). Five subjects were excluded from the analysis: Data from three participants were not analyzed due to technical issues during recording (high electrode impedance preventing data collection for safety reasons), and two participants were excluded as they could not perform the first-order task fast enough. The sample size was predefined based on power analyses conducted on pilot data, leading to a power of 0.88 (95% CI = 0.80, 0.94) with a sample size of 25 participants.

## METHOD DETAILS

### Experimental paradigm

All stimuli were prepared and presented using Python 2.7. Each trial started with the display of a 4° by 4° fixation cross presented for 500 to 1500 ms (uniform random distribution, optimized apriori to maximize design efficiency see Friston et al., 1999). Then two square boxes (size 4° by 4°) situated on each side of the fixation cross (center-to-center eccentricity of 8°) were flashed for 60 ms. In total, the two boxes contained 100 dots (diameter 0.4°) distributed unequally among them. Boxes and dots were displayed at maximum contrast on a black background. In the active condition, participants were asked to indicate which box contained most dots by pressing a key in less than 500 ms (first-order task). Responses slower than 500 ms were discouraged by playing a loud alarm sound. In the observation condition, participants were instructed to observe the image of a hand (6° by 6°) performing the first-order task by appearing on the side of the screen corresponding to one of the two boxes. They were told that the hand was controlled by a computer performing at about the same level as them to discriminate the box containing most dots. Responses in the observed condition corresponded to those in the active condition in a shuffled order, so that accuracy and response times were kept constant across conditions (see below). After the first-order response (button press or visual hand onset), a mask composed of two boxes filled with 100 dots each appeared in order to interrupt perceptual processing and ensure that the two conditions were similar in terms of visual input. After a period of time corresponding to 2 s from stimulus onset, a visual analog scale appeared instead of the mask, and participants were asked to use it to report how confident they were about their own first-order response (active condition), or about the observed first-order response (observation condition). The scale was shown for 6.5 seconds, with marks at 0 (certainty that the first-order response was erroneous), 0.5 (unsure about the first-order response) and 1.0 (certainty that the first-order response was correct). A cursor moved back and forth along the scale at slow speed (3 °/s), and participants had to press the left button at any moment when the cursor was at their chosen confidence level. The initial position and direction of the cursor was randomized and always passed through each position of the scale at least twice so that participants had one more chance were they to miss the first pass of the cursor.

Each experimental run was divided into four blocks of 12 trials, alternating between active and observation blocks. Each run started with an active block, and first-order responses in that block were shuffled and replayed in the following observation block. Importantly, the relation between response times, choice, and perceptual evidence was kept, as we shuffled trial order only. The experiment comprised six experimental runs, totalizing 144 trials per condition. During the active condition, the task difficulty was adjusted by an automatic one-up two-down staircase procedure to make the first-order performance rate converge to 71% (Levitt, 1971). The perceptual difficulty (defined as the difference in the number of dots between the two boxes) was decreased by one after one incorrect response and increased by one after two consecutive correct responses. The perceptual difficulty was pre-tuned to individual perceptual abilities by performing 96 trials of the active condition without confidence ratings prior to entering the scanner.

### Data collection

EEG data were recorded at 5000 Hz using a 63 channel setup (BrainAmp DC-amplifier, BrainProducts GmbH, Munich, Germany) synchronized to the scanner’s internal clock. Impedances of all channels were kept below 10K Ohms before entering the scanner. BOLD signal was recorded in a 3T Prisma Siemens scanner with a 32-channel coil. We used an EPI sequence (TR = 1280 ms, TE = 31 ms, FA = 64°) with 4x multiband acceleration. We acquired 64 slices of 2 x 2 x 2 mm voxels without gap (FOV = 215 mm) with slice orientation tilted 25° backward relative to the AC-PC line so as to include the cerebellum. Structural T1-weighted images were acquired using a MPRAGE sequence (TR = 2300 ms, TE = 2.32 ms, FA = 8°) with 0.9 x 0.9 x 0.9 mm voxels (FOV = 240 mm).

## QUANTIFICATION AND STATISTICAL ANALYSIS

### Behavioral analysis

Trials in which no first-order (2.0 %) or second-order response (2.9 %) was provided were excluded. Response times (RT) were defined as the time elapsed between stimulus onset and response button press (active condition), or onset of the visual hand (observation condition). Trials with RT smaller than 200 ms or higher than 500 ms (due to the loud sound) were also excluded from further analysis (13.1 %). Finally, trials from the observation condition during which the participant mistakenly pressed the response button were also excluded (12.6 %). As the exclusion criteria are not mutually exclusive, this resulted in a final number of trials of 119±5 trials in the active condition and 118±5 trials in the observation condition, out of 144 possible trials.

All continuous variables were analyzed using mixed effects models, using the lme4 (Bates et al., 2014) and lmerTest (Kuznetsova et al., 2017) packages in R. Inclusion of random effects was guided by model comparison and selection based on maximum likelihood ratio tests. The significance of fixed effects was estimated using Satterthwaite’s approximation for degrees of freedom of F statistics (Luke 2017). All statistical tests were two-tailed. Metacognitive performance was modeled using mixed effects logistic regression between first-order accuracy and confidence, with random intercept for participants and random slope for confidence. The slope of the model was interpreted as a metric for metacognitive performance (i.e., capacity to adjust confidence based on first-order accuracy). We chose this framework to analyze confidence as it is agnostic regarding the signals used to compute confidence estimates (i.e., decisional compared to post-decisional locus, see Yeung & Summerfield, 2015; Pleskac & Busemeyer, 2011), and the mixed model framework allows analyzing raw confidence ratings even if they are unbalanced (e.g., in case participants do not use all possible ratings).

### fMRI pre-processing and analysis

The functional scans were realigned, resliced and normalized to MNI space using the flow fields obtained by diffeomorphic anatomical registration through exponential linear algebra (DARTEL; Ashburner 2007). The normalized scans were smoothed using a Gaussian kernel of 5 mm full-width at half maximum (FWHM). The pre-processing was done using SPM12. We modeled the BOLD signal using a general linear model (GLM) with two separate regressors (stick functions at stimulus onset) for the active and observation condition as well as their spatial and temporal derivatives. We then parametrically modulated the regressors with three behavioral variables : the confidence ratings, the response times, and the numerosity difference between the two array of dots (i.e., perceptual evidence). Bad trials as defined in the behavioral analysis section were modeled by two separate regressors (one for active and one for observation) and their spatial and temporal derivatives. We added six realignments parameters as regressors of no interest. All second-level (group-level) results are reported at a significance-level of p < 0.05 using cluster-extent family-wise error (FWE) correction with a voxel-height threshold of p < 0.001. We used the anatomical automatic labelling (AAL) atlas for brain parcellation (Tzourio-Mazoyer et al., 2002).

### EEG pre-processing

MR-gradient artifacts were removed using sliding window average template subtraction (Allen et al., 2000). TP10 electrode on the right mastoid was used to detect heartbeats for ballistocardiogram artifact (BCG) removal using a semi-automatic procedure in BrainVision Analyzer 2. Data were then filtered using a Butterworth, 4th order zero-phase (two-pass) bandpass filter between 1 and 10 Hz, epoched [−0.2, 0.6 s] around the response onset (i.e. the button press in the active condition or the appearance of the virtual hand for in observation condition), re-referenced to a common average, and input to independent component analysis (ICA; Makeig et al., 1996) to remove residual BCG and ocular artifacts. In order to ensure numerical stability when estimating the independent components, we retained 99% of the variance from the electrode space, leading to an average of 19 (SD = 6) components estimated for each participant and condition. Independent components (ICs) were then fitted with a dipolar source localization method (Delorme et al., 2012). ICs whose dipole lied outside the brain, or resembled muscular or ocular artifacts were eliminated. A total of 8 (SD = 3) components were finally kept. All preprocessing steps were performed using EEGLAB and in house scripts under Matlab (The MathWorks, Inc., Natick, Massachusetts, United States).

### EEG univariate analysis

EEG evoked potentials were analyzed at the single trial level using a mixed effect linear regression for each channel and time point. Each model included confidence or uncertainty as dependent variables, with first-order response times and perceptual evidence (i.e., the difference in number of dots between the right and left side of the screen) as fixed effects, and a random intercept by subject. The significance of fixed effects was estimated using Satterthwaite’s approximation for degrees of freedom of F statistics, with family-wise error correction for multiple comparisons. No random slopes were added to avoid convergence failures. All analyses were performed using the tidyverse (Wickham 2017) and eegUtils (Craddock, 2018) environment in R (R core team 2018).

### EEG multivariate analysis

We derived a low dimensional description of the electrophysiological correlates of confidence using multivariate pattern analysis on single-trials. We built independent linear models in the temporal domain for each single sample within the epochs’ windows, with all the independent components retained as features. The models were evaluated using leave-one-out cross validation to avoid overfitting, and goodness-of-fit was measured by R^2^. The leave-one-out cross-validation models were also used to define the time point of maximum decoding capability within two time windows of interest ([50-200] and [200-450] ms post response). Once this time point was obtained for each window and participant, the respective EEG values estimated from the linear regressor were fed to an EEG-fMRI informed analysis (see next section).

Chance-level for decoding performance was computed using permutation statistics corrected for multiple comparisons, by repeating the whole evaluation process 1000 times while shuffling confidence rating across trials. An empirical, corrected, distribution of the null hypothesis under which R^2^ was not significantly different from zero was built by taking, for each permutation, the maximum and minimum statistics of the R^2^ throughout the whole epoch window evaluated. The corrected measure of chance level was then estimated based on the desired confidence of this distribution (fixed at *α*= 0.05).

### EEG informed fMRI analysis

To find brain-regions coactivated with decoded confidence, we built a second GLM consisting of two stick function (one for each condition), parametrically modulated by four variables; the output of the EEG confidence decoder at two time points post-response corresponding to peak R^2^ confidence decoding during the early (50 ms - 200 ms) and late (200 ms - 450 ms) time windows, the response time and the numerosity difference of the trial. We verified that empirical cross-correlation between regressors was low: rmax = 0.27 ± 0.05 and rmax = 0.22 ± 0.04 for the active and observation conditions. Excluded trials as defined in the behavioral analysis section were modeled by two separate regressors (one for active and one for observation) and their spatial and temporal derivatives. We added six realignments parameters as regressors of no interest. All second-level (group-level) results are reported at a significance-level of p < 0.05 using cluster-extent family-wise error (FWE) correction with a voxel-height threshold of p < 0.001. We used the anatomical automatic labelling (AAL) atlas for brain parcellation (Tzourio-Mazoyer et al., 2002).

### Behavioral modeling

Our models of confidence build upon a race accumulator model predicting first-order response times and choice accuracy; for every time point t (sampled at a frequency of 1000 Hz), each accumulator corresponded to the cumulative sum of independent draws from a normal distribution with unit variance and mean equal to the drift rate (v and −v for congruent and incongruent choices). The decision bound was modeled as a fixed threshold B. Non decision times were modeled by a normal distribution with mean *tnd* and standard deviation *tnd_std*. To model early errors, we added starting point variability; we allowed each accumulator to start in a non-zero state, uniformly distributed between 0 and *zvar* time the decision bound B (Purcell & Kiani, 2016).

At each iteration of the optimization procedure (see below), we generated N=1000 surrogate trials consisting in the state of the two accumulators over time and corresponding choice and RT. All parameters were fitted for the active condition, through a Nelder-Mead simplex log-likelihood minimization, comparing observed and simulated distribution of response times with a Kolmogorov-Smirnov test. To separate correct and error trials, the sign of RT was inverted for error trials. We constrained the parameters to positive values by applying an exponential transformation of the variables f(x) = exp(x), except for non-decision time and non-decision time variability which were constrained to [0,1] s by a sigmoid transformation f(x) = 1/(1+exp(-x)).

As the state of the evidence accumulation is unconstrained, we used a second stage fitting procedure to map these values to the 0-1 confidence scale. For the active condition, we sampled evidence for confidence as the state of the winning accumulator at a latency corresponding to peak performance in EEG decoded confidence. We divided the non-decision time into a sensory and an 80ms motor component (Resulaj et al.,2008). We assumed that if EEG predicted confidence best around 320 ms after the RT, then confidence would depend on the state of the accumulators 320 + 80 = 400 ms after the choice. To map the evidence to a 0 - 1 confidence scale, we used a sigmoid function:

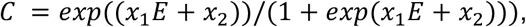

With C the resulting simulated confidence, E the accumulated evidence and x_1, x_2 two free parameters corresponding to the sensitivity and the bias of the mapping.

For the observation condition, we assumed that confidence was readout from an identical evidence accumulation process, albeit disconnected from the computer’s decisions (and response times). We thus simulated an additional 1000 surrogate trials for the observation condition but time-locked the post-decisional readout of confidence to the shuffled RTs from the active condition. The confidence readout was based on the accumulator with highest value, thus assuming a covert decision at the time of the read-out. We then fitted the parameters of the mapping as in the active condition but inversing confidence (c’ = 1-c) when the chosen accumulator deferred from the computer’s decision.

## Acknowledgements

O.B. is supported by the Bertarelli Foundation, the Swiss National Science Foundation, and the European Science Foundation. D.V. is supported by the Bertarelli Foundation and the Swiss National Science Foundation. The authors are grateful to Roberto Martuzzi, Loan Mattera, Gwénaël Birot and Gisong Kim for their help during data acquisition. We thank Elisa Filevich and Roy Salomon for constructive comments on the manuscript.

## Author Contributions

MP, NF, II developed the study concept and contributed to the study design. Testing and data collection were performed by MP, NF, II, AD, LS, SM, and MW. MP, NF, II and LS performed the data analysis. MP performed modeling work. MP, NF and II drafted the paper; all authors provided critical revisions and approved the final version of the paper for submission.

The authors declare no competing interests.

## References

Baird, B., Smallwood, J., Gorgolewski, K. J. & Margulies, D. S. Medial and lateral networks in anterior prefrontal cortex support metacognitive ability for memory and perception. J. Neurosci. 33:16657–16665 (2013).

Bang, D. & Fleming, S. M. Distinct encoding of decision confidence in human medial prefrontal cortex. Proc. Natl. Acad. Sci. 115:6082–6087 (2018).

Bogacz, R., Brown, E., Moehlis, J., Holmes, P. & Cohen, J.D. The physics of optimal decision making: A formal analysis of models of performance in two-alternative forced-choice tasks. Psychol. Rev. 113:700–765 (2006).

Boldt, A. & Yeung, N. Shared neural markers of decision confidence and error detection. J. Neurosci. 35:3478–3484 (2015).

Bonini, F., Burle, B., Liégeois-Chauvel, C., Régis, J., Chauvel, P., Vidal, F. Action monitoring and medial frontal cortex: Leading role of supplementary motor area. Science 343:888–91 (2014).

Britz, J., Van De Ville, D., & Michel, C. M. BOLD correlates of EEG topography reveal rapid resting-state network dynamics. Neuroimage 52(4), 1162–1170 (2010).

Carter, C. S., Braver, T. S., Barch, D. M., Botvinick, M. M., Noll, D. & Cohen, J. D. Anterior cingulate cortex, error detection, and the online monitoring of performance. Science 280:747 (1998).

Charles, L., Opstal, F., Van Marti, S. & Dehaene, S. Distinct brain mechanisms for conscious versus subliminal error detection. Neuroimage 73:80–94 (2013).

Debener, S., et al. Trial-by-trial coupling of concurrent electroencephalogram and functional magnetic resonance imaging identifies the dynamics of performance monitoring. J. Neurosci. 25(50):11730–11737 (2005).

Dehaene, S., Posner, M. I. & Tucker. D. M. Localization of a neural system for error detection and compensation. Psychol. Sci. 5:303–305 (1994).

Faivre, N., Filevich, E., Solovey, G., Kühn, S. & Blanke, O. Behavioural, modeling, and electrophysiological evidence for supramodality in human metacognition. J. Neurosci. 38:0322–17 (2018).

Falkenstein, M., Hohnsbein, J., Hoormann, J., Blanke, L. Effects of crossmodal divided attention on late ERP components. II. Error processing in choice reaction tasks. Electroencephalogr. Clin. Neurophysiol. 78:447–455 (1991).

Filevich, E., Forlim, C. G., Fehrman, C., Forster, C., Paulus, M., Shing, Y. L., & Kuehn, S. I know that I don’t know: Structural and functional connectivity underlying meta-ignorance in pre-schoolers. Preprint at https://www.biorxiv.org/content/early/2018/10/22/450346 (2018).

Fleck, M. S., Daselaar, S. M., Dobbins, I. G., & Cabeza, R. Role of prefrontal and anterior cingulate regions in decision-making processes shared by memory and nonmemory tasks. Cereb. Cortex 16(11):1623–1630 (2005).

Fleming, S.M., Weil, R.S., Nagy, Z., Dolan, R.J. & Rees, G. Relating introspective accuracy to individual differences in brain structure. Science 329:1541–1543 (2010).

Fleming, S.M., Dolan, R.J. The neural basis of metacognition. Philos. Trans. R. Soc. B. Biol. Sci. 367:1338–1349 (2012).

Fleming, S.M., Huijgen, J & Dolan, R.J. Prefrontal contributions to metacognition in perceptual decision making. J. Neurosci. 32:6117–6125 (2012).

Fleming, S. M., Maniscalco, B., Ko, Y., Amendi, N., Ro, T. & Lau, H. Action-specific disruption of perceptual confidence. Psychol. Sci. 26:89–98 (2015).

Fleming, S. M., & Daw, N. D. Self-evaluation of decision-making: A general Bayesian framework for metacognitive computation. Psychological review, 124(1):91 (2017).

Fleming, S. M., Putten, E. J., & Daw, N. D. Neural mediators of changes of mind about perceptual decisions. Nature. neuroscience. 21:617–624 (2018).

Gajdos, T., Fleming, S., Garcia, M. S., Weindel, G. & Davranche, K. Revealing subthreshold motor contributions to perceptual confidence. Preprint at https://www.biorxiv.org/content/early/2018/05/25/330605 (2018)

Gehring, W., Goss, B. & Coles, M. A neural system for error detection and compensation. Psychol. Sci. 4:385–390 (1993).

Gherman, S., Philiastides, M. G. Human VMPFC encodes early signatures of confidence in perceptual decisions. Elife 7:1–28 (2018).

Grimaldi, P., Lau, H. & Basso, M. A. There are things that we know that we know, and there are things that we do not know we do not know: Confidence in decision-making. Neurosci. Biobehav. Rev. 55:88–97 (2015)

Gold, J. I. & Shadlen, M. N. The neural basis of decision making. Annu. Rev. Neurosci. 30:535–574 (2007).

Hayes, S. M., Buchler, N., Stokes, J., Kragel, J. & Cabeza, R. Neural correlates of confidence during item recognition and source memory retrieval: evidence for both dual-process and strength memory theories. Journal of Cognitive Neuroscience 23(12):3959–3971 (2011).

Hebart, M. N., Schriever, Y., Donner, T. H. & Haynes, J. D. The relationship between perceptual decision variables and confidence in the human brain. Cereb. Cortex 26:118–130 (2016).

Heereman, J., Walter, H. & Heekeren, H. R. A task-independent neural representation of subjective certainty in visual perception. Front. Hum. Neurosci. 9:1–12 (2015).

Holroyd, C. B. & Coles, M. G. H. The neural basis of human error processing: Reinforcement learning, dopamine, and the error-related negativity. Psychol. Rev. 109:679–709 (2002).

Iturrate, I., Chavarriaga, R., Montesano, L., Minguez, J. & Millán, J.d.R. Teaching brain-machine interfaces as an alternative paradigm to neuroprosthetics control. Sci. Rep. 5:13893 (2015).

Kepecs, A., Uchida, N., Zariwala, H. A. & Mainen, Z. F. Neural correlates, computation and behavioural impact of decision confidence. Nature 455:227–231 (2008).

Kiani, R. & Shadlen, M. N. Representation of confidence associated with a decision by neurons in the parietal cortex. Science 324:759–764 (2009).

Kiani, R., Corthell, L., & Shadlen, M. N. Choice certainty is informed by both evidence and decision time. Neuron 84(6):1329–1342 (2014).

Kim, H. & Cabeza, R. Trusting our memories: Dissociating the neural correlates of confidence in veridical versus illusory memories. J. Neurosci. 27(45):12190–12197 (2007).

Kim, H. & Cabeza, R. Common and specific brain regions in high-versus low-confidence recognition memory. Brain research 1282:103–113 (2009).

Koriat, A. Metacognition and consciousness In: The Cambridge Handbook of Consciousness, 289–326 (2006).

Kvam, P. D., Pleskac, T. J., Yu, S. & Busemeyer, J. R. Interference effects of choice on confidence: Quantum characteristics of evidence accumulation. Proc. Natl. Acad. Sci. 112:10645–10650 (2015).

Maniscalco, B. & Lau, H. The signal processing architecture underlying subjective reports of sensory awareness. Neurosci. Conscious. 1:1–17 (2016).

Morales, J., Lau, H. & Fleming, S.M. Domain-general and domain-specific patterns of activity supporting metacognition in human prefrontal cortex. J. Neurosci. 38:2360–17 (2018).

Moritz, S., Gläscher, J., Sommer, T., Büchel, C. & Braus, D. F. Neural correlates of memory confidence. Neuroimage 33(4):1188–1193 (2006).

Murphy, P. R., Robertson, I. H., Harty, S. & O’Connell, R. G. Neural evidence accumulation persists after choice to inform metacognitive judgments. Elife 4:1–23 (2015).

O’Connell, R. G., Dockree, P. M. & Kelly, S.P. A supramodal accumulation-to-bound signal that determines perceptual decisions in humans. Nat. Neurosci. 15(12):1729–35 (2012)

Pereira, M., Sobolewski, A. & Millán, J.d.R. Action monitoring cortical activity coupled to sub-movements. eNeuro 4:1–12 (2017).

Peters, M. A. K. et al. Perceptual confidence neglects decision-incongruent evidence in the brain. Nat. Hum. Behav. 1:1–8 (2018).

Pleskac, T. J. & Busemeyer, J. R. Two-stage dynamic signal detection: A theory of choice, decision time, and confidence. Psychol. Rev. 117:864 (2010).

Pouget, A., Drugowitsch & J., Kepecs, A. Confidence and certainty: Distinct probabilistic quantities for different goals. Nat. Neurosci. 19:366–374 (2016).

Rahnev, D., Koizumi, A., McCurdy, L.Y., D’Esposito, M., Lau, H. Confidence Leak in Perceptual Decision Making. Psychol Sci 26:1664–1680 (2015).

Rahnev, D., Nee, D. E., Riddle, J., Larson, A.S. & D’Esposito, M. Causal evidence for frontal cortex organization for perceptual decision making. Proc. Natl. Acad. Sci. 113(21):6059–6064 (2016).

Rouault, M., McWilliams, A., Allen, M. & Fleming, S. M. Human metacognition across domains: insights from individual differences and neuroimaging. Personality Neuroscience 1(e17):1–13 (2018).

Shekhar, M., Rahnev, D. Distinguishing the Roles of Dorsolateral and Anterior PFC in Visual Metacognition. J. Neurosci. 38:5078–5087 (2018).

Siedlecka, M., Paulewicz, B. & Wierzchoń, M. But I was so sure! Metacognitive judgments are less accurate given prospectively than retrospectively. Front. Psychol. 7:1–8 (2016).

Ullsperger, M., Danielmeier, C. & Jocham, G. Neurophysiology of performance monitoring and adaptive behavior. Physiol. Rev. 94:35–79 (2014).

Vaccaro, A.G., Fleming, S.M. Thinking about thinking: A coordinate-based meta-analysis of neuroimaging studies of metacognitive judgements. Brain Neurosci Adv 2:1–14 (2018).

Van Den Berg, et al., A common mechanism underlies changes of mind about decisions and confidence. Elife 5:1–21 (2016).

Van Schie, H. T., Mars, R. B., Coles, M. G. H. & Bekkering, H. Modulation of activity in medial frontal and motor cortices during error observation. Nat. Neurosci. 7:549–54 (2004).

Vocat, R., Pourtois, G. & Vuilleumier, P. Parametric modulation of error-related ERP components by the magnitude of visuo-motor mismatch. Neuropsychologia 49:360–367 (2011).

Yeung, N., & Summerfield, C. Metacognition in human decision-making: Confidence and error monitoring. Phil. Trans. R. Soc. B, 367(1594), 1310–1321 (2012).

Zylberberg, A., Barttfeld, P., Sigman, M. The construction of confidence in a perceptual decision. Front. Integr. Neurosc.i 6:1–10 (2012).

## Additional references

Allen, P.J., Josephs, O. & Turner, R. A. Method for removing imaging artifact from continuous EEG recorded during functional MRI. Neuroimage 239:230–239 (2000).

Ashburner, J. A fast diffeomorphic image registration algorithm. Neuroimage 38:95–113 (2007).

Bates, D., Maechler, M. Bolker, B. & Walker, S. lme4: Linear mixed-effects models using Eigen and S4. R package version 1(7), 1–23 (2014).

Craddock, M. eegUtils: A collection of utilities for EEG analysis. R package version 0.1.13. https://github.com/craddm/eegUtils (2018)

Delorme, A., Palmer, J., Onton, J., Oostenveld, R., & Makeig, S. Independent EEG sources are dipolar. PloS one, 7(2), e30135 (2012).

Friston, K. J., Zarahn, E. O. R. N. A., Josephs, O., Henson, R. N., & Dale, A. M. Stochastic designs in event-related fMRI. Neuroimage, 10(5):607–619 (1999).

Kuznetsova, A., Brockhoff, P. B., & Christensen, R. H. B. lmerTest package: tests in linear mixed effects models. Journal of Statistical Software 82(13) (2017).

Levitt, H. C. C. H. Transformed up-down methods in psychoacoustics. The Journal of the Acoustical society of America 49(2B): 467–477 (1971).

Luke, S. G. Evaluating significance in linear mixed-effects models in R. Behavior Research Methods 49(4), 1494–1502 (2017).

Makeig, S., Bell, A. J., Jung, T. P., & Sejnowski, T. J. Independent component analysis of electroencephalographic data. Advances in neural information processing systems pp. 145–151 (1996).

Purcell, B. A. & Kiani, R. (2016). Neural mechanisms of post-error adjustments of decision policy in parietal cortex. Neuron, 89(3), 658–671.

R Core Team. R: A language and environment for statistical computing. R Foundation for Statistical Computing, Vienna, Austria. URL https://www.R-project.org/ (2018).

Resulaj, A., Kiani, R., Wolpert, D. M., & Shadlen, M. N. Changes of mind in decision-making. Nature, 461(7261), 263 (2009).

Tzourio-Mazoyer, et al., Automated anatomical labeling of activations in SPM using a macroscopic anatomical parcellation of the MNI MRI single-subject brain. Neuroimage 15(1), 273–289 (2002).

Vickers D. Decision Processes in Visual Perception. New York, NY: Academic Press (1979).

Wickham, H. tidyverse: Easily Install and Load the ‘Tidyverse’. R package version 1.2.1. https://CRAN.R-project.org/package=tidyverse (2017)

