## Supplementary material for "Disentangling the origins of confidence in speeded perceptual judgments through multimodal imaging"

### Supplementary Information

#### Supplementary behavioral analysis

We replicated the results based on mixed effects logistic regression described in the main text using second-order signal detection theory. Namely, we computed the receiver operating characteristic curve (ROC) which determines the rate of correct and incorrect responses at given confidence levels (0-25, 26-50, 51-75, 76-100%). The area between the ROC and major diagonal was taken as an index of metacognitive accuracy, which was higher in the the active (0.92 ± 0.02) compared to observation condition (0.90 ± 0.03; Wilcoxon signed rank test: $V = 163$, $p = 0.03$), in line with the interaction between accuracy and condition we found using a mixed effects logistic regression. In addition, we defined confidence bias as the log ratio between the lower and upper area separated by the minor diagonal (Kornbrot, 2006), which showed no significant difference across conditions (0.36 ± 0.14 vs. 0.27 ± 0.13; $V = 135$, $p = 0.28$). As a simple index of error awareness, we computed first-order accuracy for the first confidence bin (0-25%), and found a trend for lower choice accuracy in the active (0.01 ± 0.01) compared to observation condition (0.05 ± 0.04; $V = 10$, $p = 0.08$).

We also assessed the extent to which confidence ratings could be explained by sustained fluctuations in attention during the active and observation conditions. For that, we fitted autoregressive models to confidence for each subject and condition given the preceding five trials (i.e., lag = 5). On average, such partial autocorrelation models accounted for a small fraction of the variance in confidence ratings (subject-averaged partial coefficient: active condition = .003 ± 0.03; observation condition = -0.02 ± 0.02).

#### Follow up behavioral experiment

We devised a follow-up behavioral experiment with the same design as described above, except for the following additional points. First, we added a condition in which participants could provide their first-order response in 1500 ms (accuracy session) instead of 500 ms (speed session). Speed and accuracy sessions were run separately with counterbalanced order across participants. Each block of the speed and accuracy sessions started with a 12 trials in the active condition, followed by another 24 trials of which 12 were in the observation condition (as described in the main text), and 12 in a new continuous condition. In the continuous condition, participants were asked to report both their first and second order responses with a single key press using a horizontal scale ranging from -100 % (certainty that the left box contained more dots) to 100% (certainty that the right box contained more dots), the middle of scale corresponding to a pure guess. Trials in the observation and continuous condition were interleaved randomly, so participants could not predict in advance whether they will have to provide a first-order response or not. In total, we had 12 participants (9 females, mean age = 26.42, SD = 3.98) performing 216 trials in each of the speed and accuracy sessions, resulting in 72 trials in the active, observation, and continuous conditions.

The goal of the follow-up experiment was twofold. First, we aimed at verifying that the changes in metacognitive performance for committed and observed decisions were not due to confounding factors between the active and observation condition. For that, we compared metacognitive performance across the active and observation conditions in the speed compared to accuracy sessions, assuming that a specific improvement in error monitoring would not occur under no time pressure to provide the first-order response. We ran a mixed effects logistic regression on first-order accuracy with confidence, condition, and session as fixed effects with random slopes added for all fixed effects. The model revealed a triple interaction (odds ratios $z = 2.20$, $p = 0.03$), underlying that the interaction between condition and confidence was replicated in the speed session, but not in the accuracy session (odds ratios $z = 0.23$, $p = 0.82$). This confirms that our behavioral results in the speed session were indeed specific to a situation in which many errors occurred due to time pressure, and were not driven by low-level confounds (e.g., demand characteristics, visual saliency). As in the main experiment, response times for speeded responses in the active condition were modulated by confidence and first-order accuracy ($F(1,9.12) = 14.95$, $p = 0.004$), while the same response times in the observation condition were modulated only by first-order accuracy ($F(1,9.40) = 6.02$, $p = 0.04$) with no interaction between first-order accuracy and confidence ($F(1,8.02) = 1.65$, $p = 0.24$). This confirms the role of committed but not observed response times for confidence in speeded tasks. In the accuracy session, the same analysis revealed that responses times were modulated only by confidence (active condition: $F(1,10.26) = 15.7$, $p = 0.003$; observation condition: $F(1,9.65) = 5.79$, $p = 0.04$) and not by accuracy nor the interaction between (all $p > 0.1$).

The second goal of this follow-up experiment was to disentangle the role of decision commitment to explain the difference in metacognitive performance following committed compared to observed decisions. For that, we estimated metacognitive performance in the continuous condition, where a first-order decision was made, but for which the corresponding motor action (i.e. commitment) could not influence confidence as first and second-order responses occurred simultaneously and not sequentially as in the active condition Thus, we expected metacognitive performance in the continuous condition to resemble that of the active condition in case decision commitment was the key factor inducing changes in metacognitive performance, or instead to resemble that of the observation condition in case first-order actions shaped confidence.

We redefined confidence in the active and observation conditions as the reported probability that the correct response was right (i.e., equal to confidence in case of a right first-order response, or 1-confidence in case of a left first-order response), equivalent to the confidence rating provided in the continuous condition (see methods). We then modeled stimulus side (i.e., more dots on the right compared to left side of screen) using a mixed effects logistic regression with transformed confidence rating and condition as fixed effects, a by subject random intercept, and random slopes for confidence. In the speed session, the model revealed that the slope between accuracy and confidence was steeper in the active compared to observation condition as found previously ($z = -2.68$, $p = 0.007$), but also steeper in the active compared to continuous condition ($z = -2.44$, $p = 0.015$). Similar differences across conditions were observed regarding the model intercept (i.e., error awareness: active compared to observation condition: $z = 2.03$, $p = 0.042$; active compared to continuous condition: $z = 2.77$, $p = 0.005$) This reveals that the fact of committing to a decision (i.e. a decision leading to a first-order action) improves subsequent confidence estimates. In the accuracy session, the only significant effect was confidence ($z = 13.02$, $p < 0.001$), with no main effect of condition or interaction between condition and confidence (all $-1 < z < 1$).

**Supplementary EEG analysis**

To disentangle the contribution of first and second order responses to electrophysiological markers of confidence, we ran the same analysis as reported in the main text, but this time modeling EEG amplitude as a function of uncertainty, defined as the absolute distance between the reported confidence and 50 % confidence (|1- |2*(confidence - 0.5)||) so that uncertainty was symmetrical for correct and incorrect first-order responses. In both conditions, effects of uncertainty were found over frontocentral electrodes, respectively 392 and 500 ms post-response ($p < 0.05$, fdr-corrected), suggesting that early correlates of confidence are mostly due to error monitoring.

We also assessed motor responses in the active and observation condition by computing lateralized readiness potentials (LRP Coles 1988). We defined LRPs for each subject as (C3r - C4r + C4l - C3) / 2. A one-sample t-test of LRP amplitude against zero was performed, followed by false-discovery rate correction. In the active condition, a deflection starting 80ms before the first-order response (SFigure 1). In the observation condition, a smaller potential was found later, 176 ms following the observed response, suggestive of a response evoked by the visual feedback.

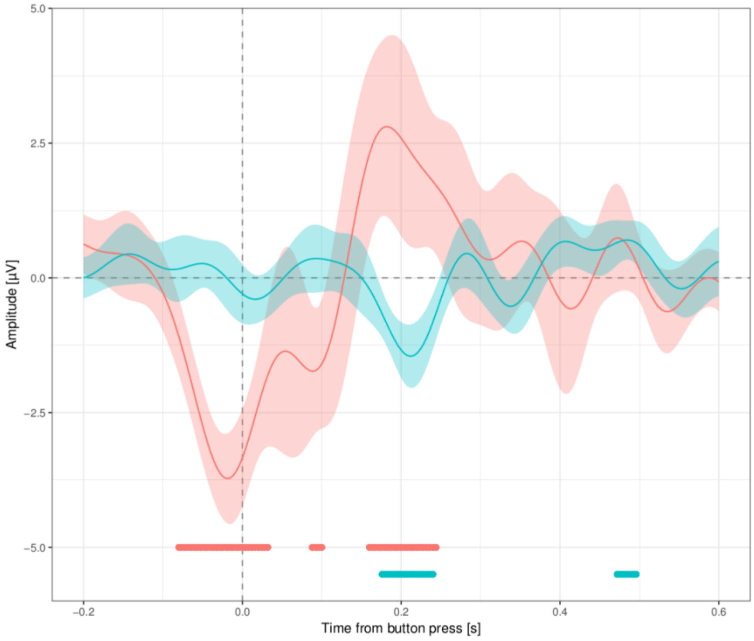


Supplementary Figure 1: Lateralized readiness potentials for committed (in red) and observed (in blue) first-order responses. The horizontal lines under each curve indicates time points for which the LRP differs from zero ($p < 0.05$ fdr-corrected).

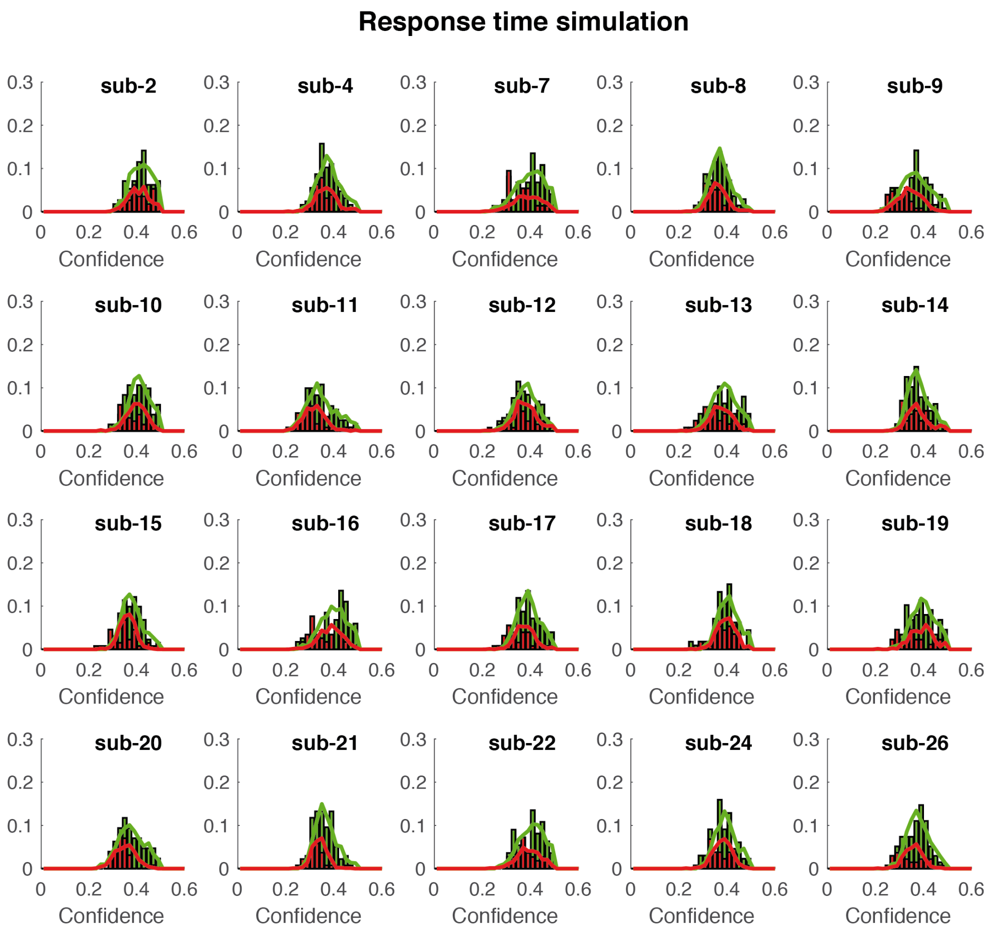


Supplementary Figure 2: Response time simulation for each subject. Error trials have their RT inverted. Histograms represent RT data (red for error and green for correct). Red and green traces represent simulated RT using a race accumulator model.

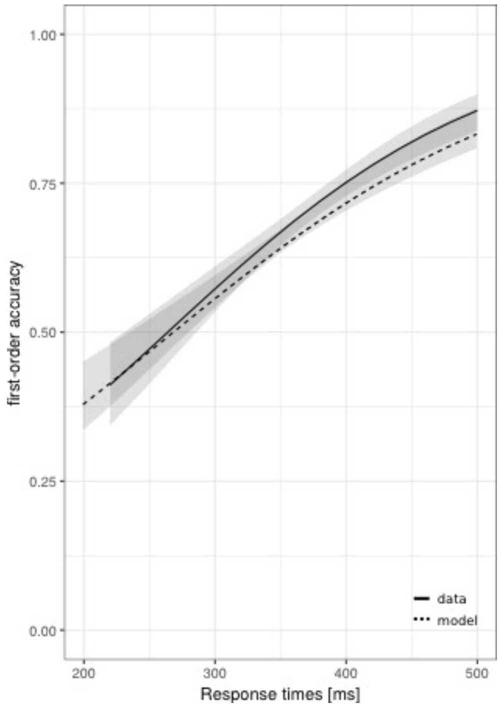

Supplementary Figure 3: Increase in choice accuracy with response time. The solid thick line is the logistic fit of the data, averaged across subjects. The dashed thick line is the logistic fit of the simulated data, averaged across subjects. Thin lines represent 95%-CI.

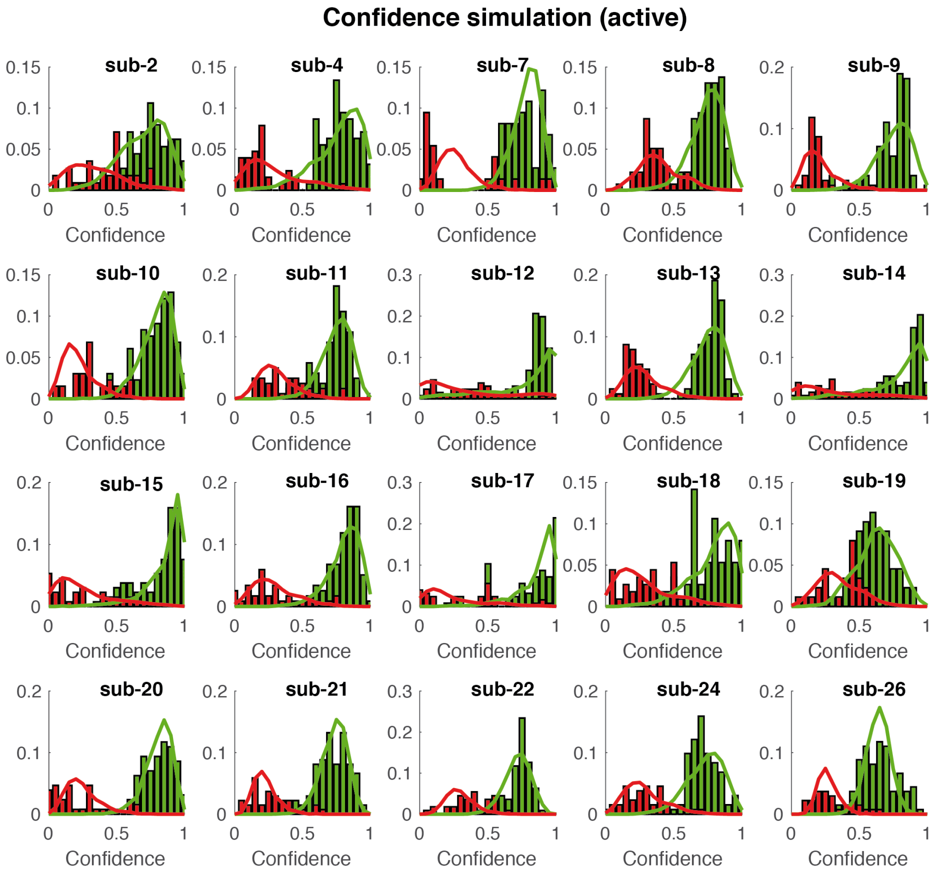


Supplementary Figure 4: Confidence simulation for the active condition for each subject. Histograms represent confidence data (red for error and green for correct). Red and green traces represent simulated confidence using a race accumulator model and a second-order model of confidence.

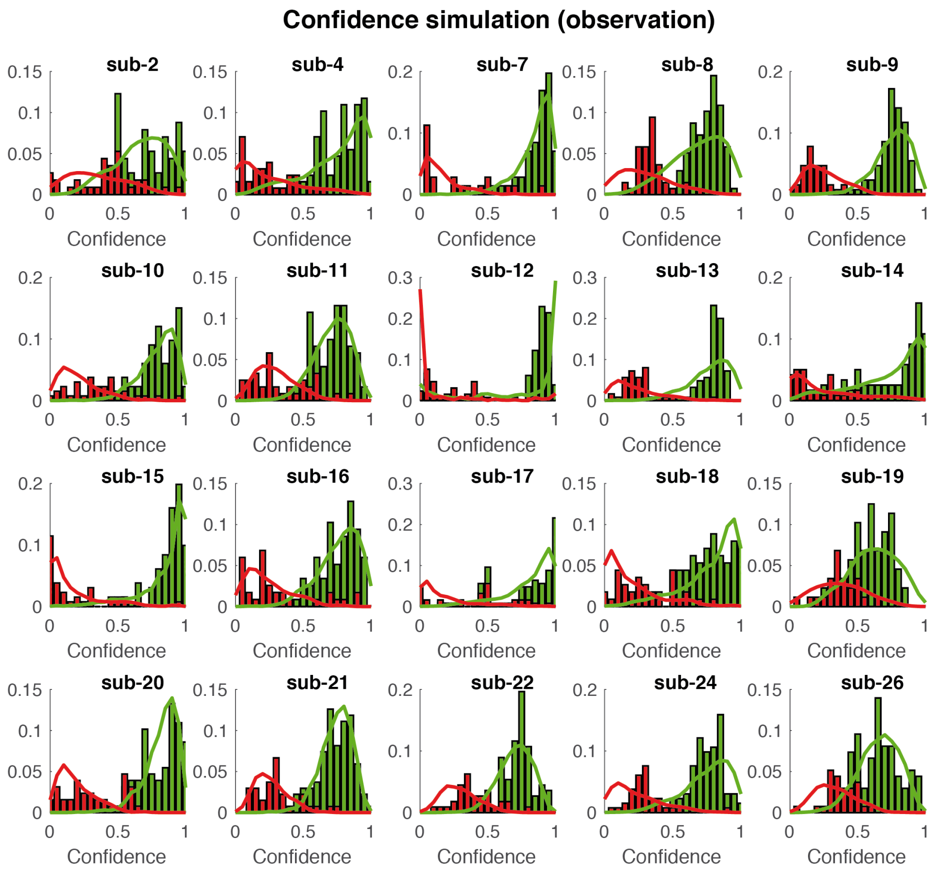


Supplementary Figure 5: Confidence simulation for the observation condition for each subject. Histograms represent confidence data (red for error and green for correct). Red and green traces represent simulated confidence using a race accumulator model and a non-decisional model of confidence.

Supplementary table 1. Significant activations for confidence (fMRI analysis)

| Condition | Label | Side | Voxels | MNI peak | T-value | p-value |
|---|---|---|---|---|---|---|
| Active > 0 | Occipital | L | 564 | -21 -98 8 | 5.07 | <0.001 |
| | Ventral striatum | L/R | 353 | 3 3 -8 | 4.84 | 0.001 |
| | Putamen | L | 336 | -24 -3 22 | 5.05 | 0.002 |
| | vmPFC | L | 322 | -14 34 -8 | 5.72 | 0.002 |
| | Occipital | R | 258 | 16 -98 3 | 4.67 | 0.009 |
| | HC | R | 181 | 33 -24 -4 | 4.85 | 0.049 |
| Active < 0 | (*) | L/R | 17572 | -6 15 64 | 8.59 | <0.001 |
| | IPL | L | 2737 | -26 .69 58 | 6.48 | <0.001 |
| | aPFC | L | 1347 | -34 51 14 | 4.90 | <0.001 |
| | IFG | R | 770 | 51 22 8 | 4.99 | <0.001 |
| | IPL | R | 409 | 36 -40 45 | 4.23 | <0.001 |

| | | | | | | |
|---|---|---|---|---|---|---|
| | MTL | L | 403 | -46 -30 -6 | 4.70 | <0.001 |
| | Precuneus | L | 266 | -12 -66 48 | 4.61 | 0.007 |
| Observation > 0 | STL | L | 2461 | -56 -33 14 | 5.82 | <0.001 |
| | STL | R | 2107 | 62 -20 16 | 6.11 | <0.001 |
| | Cuneus | L | 2050 | -4 -82 30 | 5.44 | <0.001 |
| | vmPFC | L/R | 1215 | 4 33 -18 | 4.79 | <0.001 |
| | HC/AM | L | 796 | -26 -4 -15 | 5.44 | <0.001 |
| | Paracentral lobule | R | 502 | 3 -34 56 | 5.81 | <0.001 |
| | Occipital | L | 399 | -21 -99 9 | 4.95 | 0.001 |
| | Postcentral | L | 383 | -30 -21 44 | 5.89 | 0.001 |
| | Precuneus | L/R | 376 | -4 -56 24 | 4.78 | 0.001 |
| | Postcentral | R | 373 | 24 -39 69 | 4.11 | 0.001 |
| | HC | R | 262 | 18 -15 -15 | 5.36 | 0.008 |
| | MTL | L | 202 | -60 -9 -16 | 5.54 | 0.03 |
| | Occipital | R | 191 | 14 -100 10 | 5.14 | 0.039 |
| Observation < 0 | (pre-)SMA | L/R | 853 | -4 21 52 | 4.69 | <0.001 |
| | IPL | L | 683 | -40 -45 45 | 4.94 | <0.001 |
| | IPL | R | 612 | 34 -46 44 | 4.50 | <0.001 |
| | Precentral | L | 381 | -50 8 45 | 4.54 | 0.001 |
| | Insula | L | 310 | -32 20 -3 | 4.45 | 0.003 |
| | Insula | R | 228 | 33 21 -6 | 4.31 | 0.017 |
| | IFG | L | 223 | -57 16 15 | 4.03 | 0.018 |

(*) bilateral supplementary motor area and dorsal anterior cingulate cortex/left superior, middle and inferior frontal cortices (SFC,MFC,IFG) and anterior insula (AI)

Supplementary table 2. Significant differences for confidence in both conditions (fMRI analysis)

| Condition | Label | Side | Voxels | MNI peak | T-value | p-value |
|---|---|---|---|---|---|---|
| Active < Observation | pMFC | L | 1244 | -14 22 64 | 7.06 | <0.001 |
| | Precentral | R | 164 | 28 -10 46 | 6.06 | 0.033 |
| | Insula | L | 199 | -40 6 -4 | 5.82 | 0.012 |
| | dACC | L/R | 181 | -3 27 38 | 4.82 | 0.021 |

Supplementary table 3. Significant conjunctions for confidence (fMRI analysis)

| Condition | Label | Side | Voxels | MNI peak | T-value | p-value |
|---|---|---|---|---|---|---|
| (Active < 0) ∩ (Observation < 0) (late) | pMFC | L/R | 606 | -4 21 52 | 4.69 | <0.001 |
| | IPL | L | 552 | -40 44 45 | 4.82 | <0.001 |
| | Precentral | L | 350 | -50 8 45 | 4.54 | 0.001 |
| | AI | L | 281 | -32 20 -3 | 4.45 | 0.005 |
| | IFG | L | 223 | -57 16 15 | 4.03 | 0.018 |

Supplementary table 4. Significant activations for decoded confidence (EEG-informed fMRI analysis)

| Condition - latency | Label | Side | Voxels | MNI peak | T-value | p-value |
|---|---|---|---|---|---|---|
| Active < 0 (early) | pMFC | L/R | 1927 | -12 12 66 | 5.65 | <0.001 |
| | IFG/AI | L | 1233 | -48 16 6 | 5.08 | <0.001 |
| | MFG | L | 165 | -38 6 52 | 5.38 | 0.025 |
| Observation > 0 (early) | vmPFC | L/R | 756 | 10 36 -2 | 4.78 | <0.001 |
| | vmPFC | L/R | 361 | 0 30 -12 | 5.37 | 0.001 |
| Active < 0 (late) | pMFC | L/R | 2689 | -21 12 60 | 5.10 | <0.001 |
| | IPL | L | 1346 | -45 -34 42 | 4.87 | <0.001 |
| | Precentral | L | 1155 | -39 0 39 | 5.02 | <0.001 |
| | AI | L | 1120 | -32 24 -3 | 5.04 | <0.001 |
| | aPFC | L | 326 | -34 40 10 | 4.01 | 0.003 |
| | IFG | L | 263 | -36 24 27 | 4.04 | 0.01 |
| | MFG | L | 262 | -36 20 42 | 4.36 | 0.01 |
| | IFG | R | 258 | 44 26 -3 | 4.11 | 0.011 |
| | pMFC | R | 225 | 8 4 68 | 4.50 | 0.022 |
| Observation < 0 (late) | IFG | L | 191 | -50 16 4 | 4.72 | 0.046 |
| Active < Observation (early) | AI | L | 266 | -32 14 -9 | 4.63 | 0.009 |
| Active < Observation (late) | aPFC | L | 248 | -28 54 9 | 4.29 | 0.013 |

#### References:

Holroyd, C. B., Dien, J., & Coles, M. G. Error-related scalp potentials elicited by hand and foot movements: evidence for an output-independent error-processing system in humans. *Neuroscience letters* **242**(2):65-68 (1998).

Kornbrot, D. E. Signal detection theory, the approach of choice: Model-based and distribution-free measures and evaluation. *Perception & Psychophysics* **68**(3), 393-414 (2006).
